# Mobilisation and analyses of publicly available SARS-CoV-2 data for pandemic responses

**DOI:** 10.1101/2023.04.19.537514

**Authors:** Nadim Rahman, Colman O’Cathail, Ahmad Zyoud, Alexey Sokolov, Bas Oude Munnink, Björn Grüning, Carla Cummins, Clara Amid, David Nieuwenhuijse, Dávid Visontai, David Yu Yuan, Dipayan Gupta, Divyae Prasad, Gábor Máté Gulyás, Gabriele Rinck, Jasmine McKinnon, Jeena Rajan, Jeff Knaggs, Jeffrey Edward Skiby, József Stéger, Judit Szarvas, Khadim Gueye, Krisztián Papp, Maarten Hoek, Manish Kumar, Marianna Ventouratou, Marie-Catherine Bouquieaux, Martin Koliba, Milena Mansurova, Muhammad Haseeb, Nathalie Worp, Peter W. Harrison, Rasko Leinonen, Ross Thorne, Sandeep Selvakumar, Sarah Hunt, Sundar Venkataraman, Suran Jayathilaka, Timothée Cezard, Wolfgang Maier, Zahra Waheed, Zamin Iqbal, Frank Møller Aarestrup, Istvan Csabai, Marion Koopmans, Tony Burdett, Guy Cochrane

**Affiliations:** European Molecular Biology Laboratory, European Bioinformatics Institute, Wellcome Genome Campus, Hinxton, Cambridgeshire, UK; Technical University of Denmark, Anker Engelunds Vej 101, 2800 Kongens Lyngby, Denmark; Eötvös Loránd University, H-1053 Budapest, Egyetem tér 1-3, Hungary; Erasmus Medical Center, Wytemaweg 80, 3015 CN Rotterdam, Netherlands; University of Freiburg, Friedrichstr. 39, 79098 Freiburg

## Abstract

The COVID-19 pandemic has seen large-scale pathogen genomic sequencing efforts, becoming part of the toolbox for surveillance and epidemic research. This resulted in an unprecedented level of data sharing to open repositories, which has actively supported the identification of SARS-CoV-2 structure, molecular interactions, mutations and variants, and facilitated vaccine development and drug reuse studies and design. The European COVID-19 Data Platform was launched to support this data sharing, and has resulted in the deposition of several million SARS-CoV-2 raw reads. In this paper we describe (1) open data sharing, (2) tools for submission, analysis, visualisation and data claiming (e.g. ORCiD), (3) the systematic analysis of these datasets, at scale via the SARS-CoV-2 Data Hubs as well as (4) lessons learned. As a component of the Platform, the SARS-CoV-2 Data Hubs enabled the extension and set up of infrastructure that we intend to use more widely in the future for pathogen surveillance and pandemic preparedness.

## Introduction

On December 27, 2019, public health authorities of Wuhan, China were notified of a small number of pneumonia-like cases, seemingly linked to visits to a market^1^. The cluster was quickly identified as being caused by a novel coronavirus, soon thereafter named Severe Acute Respiratory Syndrome Coronavirus 2 (SARS-CoV-2) due to its genetic similarity to SARS-CoV, the virus causing the SARS outbreak in 2003^1^. Despite initial optimism that the outbreak could be controlled by enforcing stringent public health measures, the virus spread to different cities in the region, and soon cases presenting with the disease (Coronavirus Disease 2019 or COVID-19) emerged on every continent. By 11th March 2020, the World Health Organisation (WHO) declared this COVID-19 outbreak a pandemic^2^, triggering a massive global response. One of the hallmarks of the public health response during this pandemic was the extensive application of pathogen genomic sequencing, which brought in scientific institutes and consortia to mobilise sequencing efforts and participate in the sharing of COVID-19 biodata^3, 4^.

The European COVID-19 Data Platform was launched in April 2020, as part of the European Molecular Biology Laboratory’s (EMBL) response to the COVID-19 pandemic. The data platform comprises three main components. First, the COVID-19 Data Portal provides an interface to access and browse a wealth of publicly available datasets, tools and resources^5^. Second, the Federated European Genome-phenome Archive (FEGA) supports the sharing of sensitive human genotypic and phenotypic data^6^. Third, the SARS-CoV-2 Data Hubs, the topic of this paper, offer a toolbox to those working with viral sequence data to support management, sharing, analysis and interpretation. A particular focus of our work with the SARS-CoV-2 Data Hubs was to make public data more easily available to open research around the world, as well as to ensure the sharing of raw sequencing data in addition to consensus sequences, along with a host of other data types (as presented within the COVID-19 Data Portal). By doing so, re-analysis can be done allowing for evaluation of published consensus sequences, using other tools for assembly, minority variants determination, etc. This involved mobilisation of teams, infrastructure and data to support worldwide scientific research around SARS-CoV-2 sequence data.

Coupling a public health emergency with the increasingly reduced sequencing costs, has resulted in a wealth of data sharing to a level never seen before and currently nearly 25% (24.88%) of the total number of raw read records in the European Nucleotide Archive (ENA) are from SARS-CoV-2. Therefore, there was a need to facilitate submission and analysis of this explosively expanding dataset. Here we describe the development of public SARS-CoV-2 Data Hubs and their components, specifically designed to facilitate further analysis by the wider research community. We describe the community-owned/driven data hub model, the submission services and tools, analytical workflows, visualisation tools and search and retrieval via the COVID-19 Data Portal.

## Overview

The SARS-CoV-2 Data Hubs (Fig.1) build on a model developed through the COMPARE project^7^. This involved the provision of protected databases, in which groups of collaborating scientists could share their pre-publication private data as well as results of analyses, prior to data release. EMBL-EBI implemented a modular system centring around data storage modules, connecting to analysis compute, and including several components and tools, described further in the COMPARE Data Hubs^7^. The public SARS-CoV-2 Data Hub described here, consists of several components, building on the components described in the COMPARE Data Hubs, including a set of submission tools, analysis workflows and infrastructure, portals or interfaces for data and metadata search and retrieval, and visualisation tools. As the data hubs sit on top of ENA^8^, within EMBL-EBI^9^ infrastructure, they enable for integration with all existing metadata and data models in ENA and the connected databases of the International Nucleotide Sequence Database Collaboration (INSDC)^10^. The data hub model was built to allow easy re-use in the future, for other diseases and datasets, facilitating long-term sustainability and open science. A specific focus has been on enabling sharing of raw read data to allow for standardised analysis of all public data, and future re-analyses using novel bioinformatic tools. The data hubs are configurable, enabling users to choose appropriate tools according to their needs. This allows users in turn to focus more on datasets held within data hubs, bringing tools to the data. The development of key components of the data hubs included collaborations with partners in the VEO project^11^: Technical University of Denmark (DTU), Eötvös Loránd University (ELTE) of Hungary and Erasmus Medical Centre (EMC) of Netherlands.

**Figure 1).**
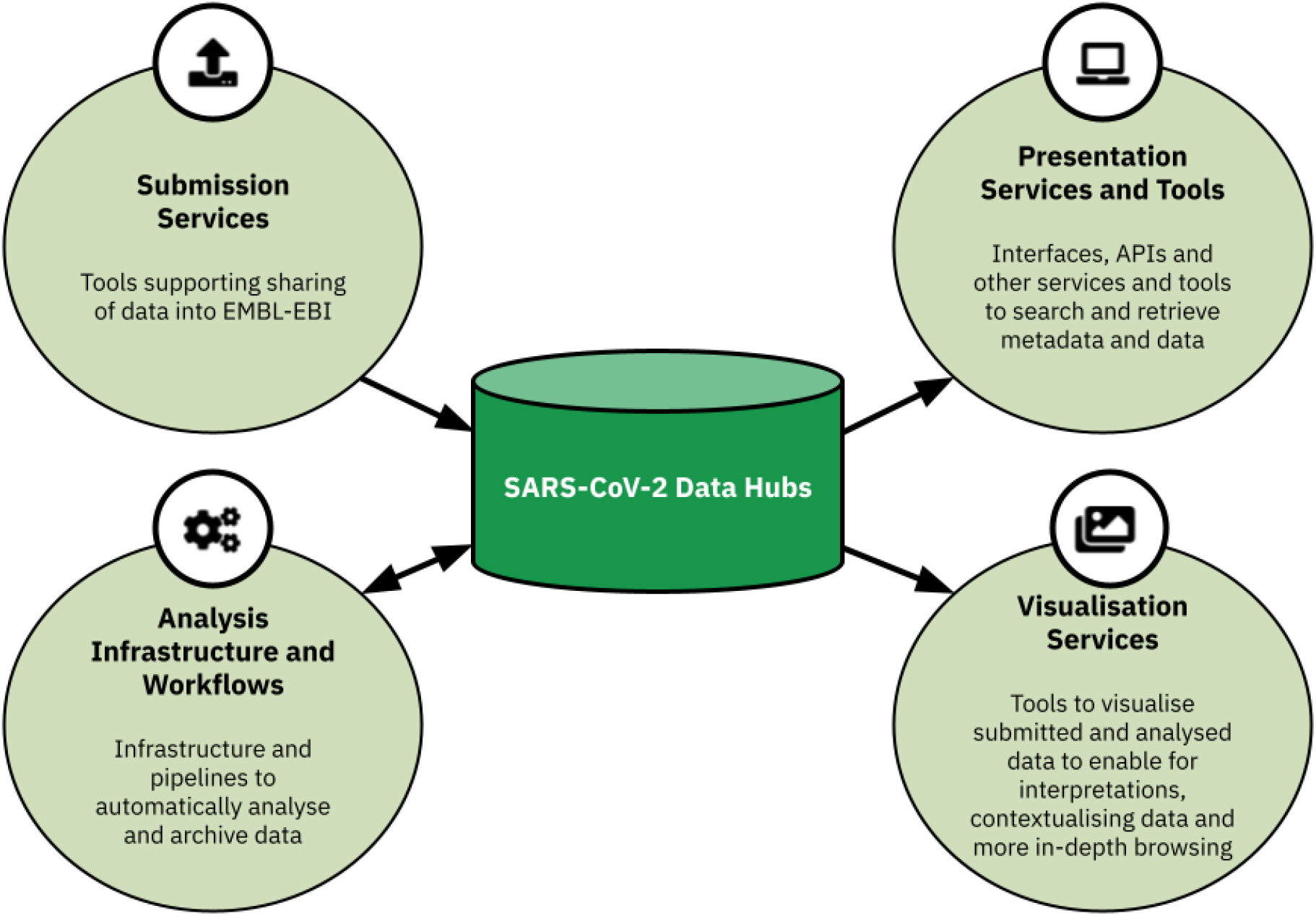
The SARS-CoV-2 Data Hubs and their components. This includes submission services, analysis infrastructure and workflows, presentation services and tools and visualisation services.

## Methods

In April 2020, European Union president Ursula von der Leyen commissioned the development of the European COVID-19 Data Platform, an initiative aiming to leverage data sharing infrastructure for research during the pandemic. The VEO consortium assembled a taskforce to help address the software-based variability among shared genomic data. As part of the data mobilisation efforts for the COVID-19 Data Portal, EMBL-EBI encouraged and supported mobilisation of raw read sequencing data from data providers, i.e. prior to any assembly and sharing of annotated genomes, as seen through (and in addition to) GISAID. Through meetings with users and data providers interested in sharing data, their challenges, feedback and requirements in handling the vast amount of data were identified, which helped drive tool development. Utilising this feedback, we developed, tested and reviewed tools to further support the mobilisation of data. Workflows, visualisation tools and components were developed for the analysis and interpretation of all mobilised data. These aspects supported the development of the public SARS-CoV-2 Data Hub, and included a focus on sustainability beyond the COVID-19 outbreak, to support re-use in any future outbreaks. This involved a large group of researchers and collaborators within the consortium, working on several different aspects, which have been described below.

### Analysis workflow for processing raw read datasets

#### Systematic Analysis

A flow for the systematic analysis of publicly submitted read data within the COVID-19 Data Portal was established, depicting an example of a public SARS-CoV-2 Data Hub. The data hub comprises several components, spanning the adaptation, development, extension and integration of workflows, an analysis management system, and data flow management within the data platform for dissemination and visualisation tools (along with their integration).

A key feature is that submitted raw sequence data are all uniformly processed through a platform specific (Illumina or Oxford Nanopore) pipeline to generate consensus sequences and to identify mutations in the submitted datasets for variant calling. By using this approach, the data produced can be compared with confidence that variation introduced through implicit biases of different analysis approaches used in different laboratories is excluded. Therefore, when combined with quality thresholds and control for sequencing platforms used, variation observed between samples^12^ most likely reflects true biological variation^13^.

### Workflows

#### COVID-19 Sequence Analysis Workflow

Public read data (runs) submitted to the ENA were processed through the COVID-19 Sequence Analysis Workflow^14^, if the run metadata specifies “ILLUMINA” as the instrument platform. The workflow first trims reads using Trimmomatic^15^ SLIDINGWINDOW:5:30 MINLEN:50. Reads are then mapped to the SARS-CoV-2 reference genome (NC_045512.2) using bwa mem^16^. Samtools^17^ mpileup -Q 30 -d 8000 is used to generate a tab separated single base pair resolution coverage file. Variants are called from indexed and mapped reads using LoFreq^18^. Two different sets of variants are produced from this process, one that is unrestricted, and a second that only contains variants with an allele frequency (AF) greater than 0.25. Variants are annotated using SnpEff^19^. A consensus sequence is generated from these variant files using a custom python script^20^.

#### Nanopore Analysis Workflow

Public raw read data (runs) submitted to the ENA were processed through the Nanopore Analysis Workflow^14^ (incorporated within the COVID-19 Sequence Analysis Workflow), if the run metadata specifies “OXFORD_NANOPORE” as the instrument platform. The workflow first trims the start and end of each read by 30 bps using cutadapt^21^. Trimmed reads are then mapped to the SARS-CoV-2 reference genome using minimap2^22^. Variants are called by registering any variant in the mapping result occurring at more than a 10% variant frequency. Below this frequency the number of clearly erroneous variants (in particular single nucleotide deletions) increased dramatically. At the same time majority voting assigns variants as “MAJOR” variant by determining if the most abundant nucleotide at each position in the genome is the same or different from the reference. All called variants are stored in the unfiltered VCF, while only the “MAJOR’’ variants are stored in the filtered VCF. The filtered VCF is in turn used to generate the consensus sequence by incorporating the variants in the reference genome and masking low coverage regions. All custom scripts to perform calling, filtering and consensus calling are available on GitHub^23^.

#### Lineage Assignment

To provide lineage annotations, the Pangolin^24^ workflow has been integrated at EMBL-EBI to assign PANGO lineages to both user submitted and systematically analysed consensus sequences. In addition, World Health Organisation (WHO) variants of concern (VOCs) and variants of interest (VOIs) assignments are included, if applicable. The Pangolin workflows^25^ integration was implemented in Snakemake^26^.

#### Representative Sequences

To support the wider scientific community in generating phylogenies or other comparative analyses, ‘representative sequences’ were identified^27^. This essentially provides a backbone of sequences, which users can retrieve, with a sequence per Pango lineage, allowing rapid assessment of placement of a newly submitted sequence. Here, sequences are selected based on a two step filtering process. First, the output of a Pangolin run on all consensus sequences archived at the ENA is parsed and sequences with a scorpio support value^24, 28^ of less than 0.95 are removed. The scorpio support value is defined as the proportion of defining variants which have the alternative allele in the sequence. This step also removes low coverage sequences as these do not get assigned a scorpio support value. Second, the earliest appearance of a unique Pangolin lineage (e.g. A.1, B.1.1.7) in this list is selected based on the “collection date” ENA metadata field, after 1st Sept 2020. This date is selected to filter out sequences with potentially erroneous collection dates that pre-date the earliest observation of a lineage. This is an automated curation process using a custom python script^29^.

### Visualisations

#### COVID-19 Phylogeny

To visualise the evolution and lineages of SARS-CoV-2 from genomic data, a phylogenetic tree of submitted SARS-CoV-2 sequence data was set up using the Evergreen phylogenetic analysis workflow^30^. The workflow^31^ employs a reference based alignment strategy, where the input sequences are matched to references by genomic similarity calculated on k-mers. The workflow processes raw reads from single molecule sequencing technologies in addition to short read technologies and consensus sequences. Hamming distances are calculated between sequences in the alignment, excluding insertions and deletions, and a neighbour-joining tree is inferred on the resulting distance matrix. Further details on the methods applied for the COVID-19 phylogeny can be found on the COVID-19 Data Portal.

PhyloDash^32^ was used to present Evergreen output to users. PhyloDash is a reusable React component connecting interactive modules of Mapbox with OpenStreetMap^33^, the phylogenetic tree with Phylocanvas.gl^34^, a metadata table, and a timeline.

#### Variant Browsing

The Kooplex collaboration platform^35^ was used to develop CoVEO^36^. This application was developed and used for variant and sample browsing from systematically analysed raw reads. To obtain details on the number of new weekly COVID-19 cases, external data from the John Hopkins Coronavirus Resource Center^37^, was required for an estimation of the percentage of new cases that have been sequenced in countries. Input samples were classified into variants of concern (VOC) or variants under investigation (VUI) groups based on key mutations in the database. Kooplex enables direct access to the database containing SARS-CoV-2 data and the system can be used for direct advanced data analysis and visualisation to answer scientific questions, for example identifying samples carrying mutations in SARS-CoV-2 PCR primer regions that may affect the testing efficiency^38^.

## Results

**Table 1).** Requirements that were identified from discussions with stakeholders and users. This also includes tools to address trends and aspects of interest for the wider scientific community at the time, as noticed from literature and social media.

| Requirement | Description | Reason for requirement | Tool(s) developed |
| --- | --- | --- | --- |
| Support a more-bulk method to submit COVID-19 genomes. | To submit SARS-CoV-2 genomes, users were restricted to using Webin-CLI. This tool, although relatively fast and generally supportive towards larger scale submissions, was not efficient in support for high-throughput submissions to the levels seen during the COVID-19 pandemic. | To support the high volume of SARS-CoV-2 genomes from public health genome sequencing programmes. | Webin SARS-CoV-2 Genome Submission Web API |
|  |  |  | ENA Bulk Webin-CLI Tool |
| Support for a simpler submissions interface. | Although interactive submissions to the ENA exist, this does require prior knowledge of the ENA submission process. | To provide a simple 'drag and drop' like tool to ease the submission process for prospective non-informatics based users. | SARS-CoV-2 Drag and Drop Uploader |
| Support large-scale analysis of SARS-CoV-2 raw read data. | To process the large amount of SARS-CoV-2 raw read data, sufficient management, scheduling, analysis, and finally submission of analysis products were required. | Appropriate scheduling, management, processing and data provenance/trail was required to adapt the process to the quantity of data. | ENA Pathogen Analysis System |
|  |  |  | ENA Analysis Submitter |
| Support data claiming and attribution | When data is shared, affiliation and information detailing the data submitter or generator is held within and presented in data records. | Support researchers in data sharing by including additional methods in acknowledgement and attribution. | ORCID Data Claiming |
|  |  |  | Dataset DOIs |
| Annotate sequencing data in-line with WHO naming | Submitted SARS-CoV-2 genomes did not include lineage information as standard. | Support researchers in greater interpretation of data, and tailored searching for datasets of interest. | Integrated Pangolin pipeline setup |
| Tools to better support users in contextualising, visualising and interpreting genomic data | For SARS-CoV-2, there were no tools integrated into the platform to support the user in interpreting their data in greater detail. | Provide tools and visualisations to support researchers and other users in their interpretations | COVID-19 Phylogeny |
|  |  |  | CoVEO |

### Submissions

#### Tools

##### Webin SARS-CoV-2 Genome Submission Web API

For large-scale programmatic submitters of SARS-CoV-2 genomic data, users mentioned that existing routes for genome submission at the ENA, whereby metadata and sequence are separated, could be more intuitive and thereby reduce any bottlenecks when preparing and submitting data. The Webin SARS-CoV-2 Genome Submission Web Application Programming Interface (API) (launched in April 2021) is a dedicated tool for the programmatic submission of SARS-CoV-2 genome assemblies. During testing, the tool was seen to halve the amount of time taken per submission, compared to an equivalent submission made using the existing method, Webin-CLI^39^, both using a single thread. Assembled sequence data and metadata are provided within the same JSON payload, and are associated to a pre-registered ENA study and sample by specifying the respective accessions. By providing a validate option, it enables testing of a submission in order to identify any errors, prior to genome submission. Here, an API addresses the bottleneck, as this can be easily incorporated into scripts and tools written in different programming languages. It expands the existing assembly submission routes offered by the ENA, and is currently utilised by several national SARS-CoV-2 data brokers, and other high volume SARS-CoV-2 data submitters.

##### SARS-CoV-2 Drag and Drop Uploader

Within the community and from feedback obtained, there was a need for the development of a simpler interface and submission process to share data to keep up with the rapid expansion of the genomic sequencing effort. The aim of this type of interface was to support sharing via another interactive route, without the need for extensive bioinformatics skills or prior knowledge of the ENA submission system. This resulted in the first new submission tool developed and launched in May 2020, the SARS-CoV-2 Drag and Drop Uploader, a browser-based ENA uploader^40^.

This tool (Fig.2) simplified the submission process at the ENA, by enabling users to drag and drop their sequence data files, accompanied by a metadata spreadsheet. Behind the scenes, data is stored on a cloud-based storage bucket, prior to being submitted to the archive and data hub. For optimal performance and user-experience, the tool is limited to around 30GB per upload, with no actual limit for the number of uploads. The speed of upload will be dependent on the submitter’s internet connectivity. To use the tool, submitters are first required to contact the ENA SARS-CoV-2 helpdesk via the form on the web page^40^ and request the following: a secure login key unique to their ENA submission account, and an appropriate metadata spreadsheet for their data type.

**Figure 2).**
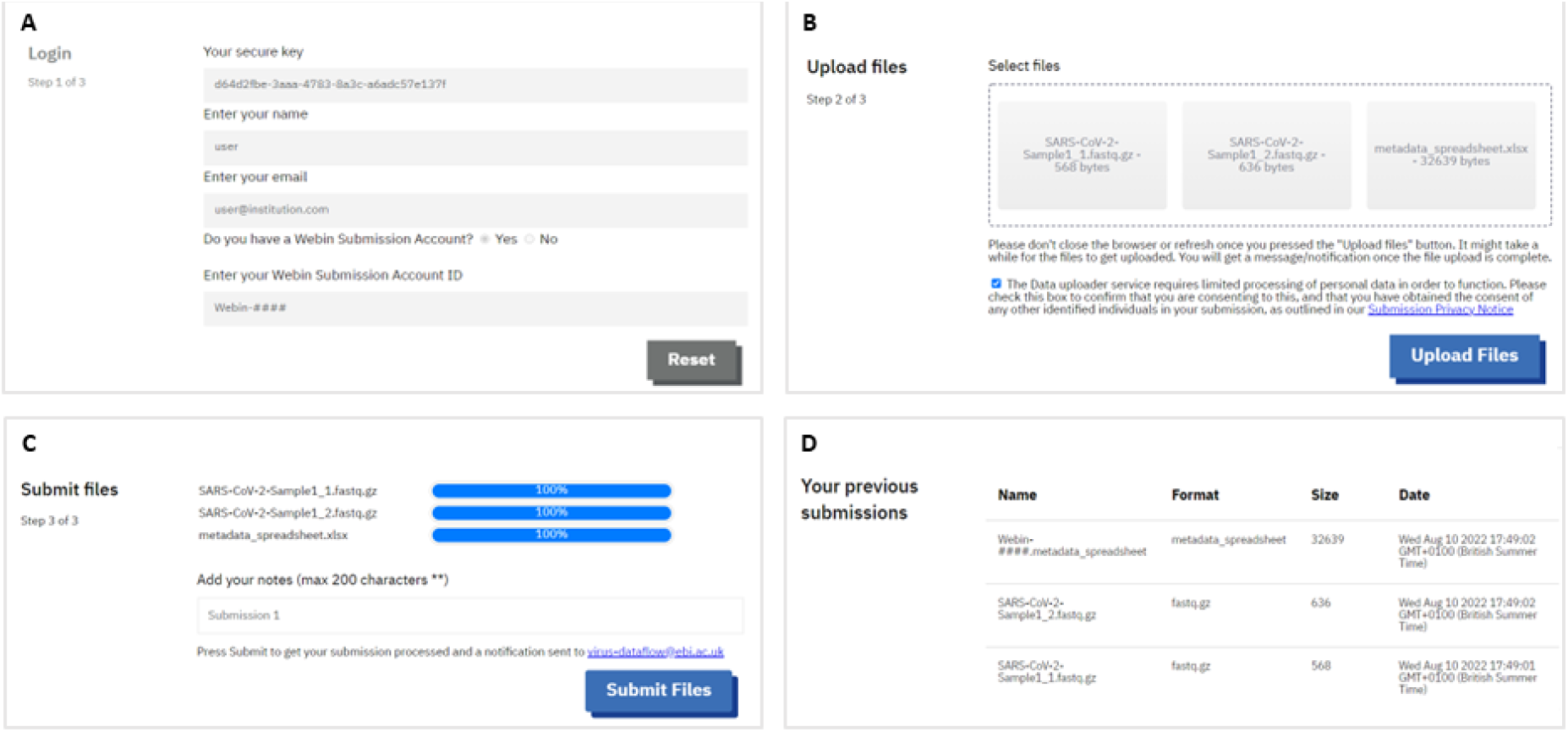
SARS-CoV-2 Drag and Drop Uploader Tool Interface. (A) Tool login page for users requiring a submission key (provided by the helpdesk team), brief contact details and their Webin Account ID. (B) Drag and drop area for users to deposit their data files and metadata spreadsheet, before uploading. (C) Upload progress area for users to monitor their file status. Submission occurs after successful upload for validation and processing at the ENA. (D) Submission history area where users can see their previous submissions through the tool.

Since its release in early 2020, there have been a total of 33 submissions with the Drag and Drop Uploader, with ∼50% of users being new submitters to ENA. 83% of these data submissions are raw reads (i.e FASTQ or BAM files) while the remaining 17% are assembled/consensus sequence submissions (mainly in FASTA format).

To support FAIR data principles, submissions to the ENA are structured via a metadata model. There was an even split between the number of end-to-end submissions made using the tool (from ENA top level project registration through to metadata and then data file submission), and part-submissions (e.g. only data file submission, whilst referencing pre-submitted project and metadata).

In a user survey with a total of 9 respondents, 6 rated the tool ‘excellent’ and 5 said they would use the tool again. Indeed 5 tool submissions were repeated across 2 users.

##### Additional Submission Tools

###### GISAID Spreadsheet Conversion Tool

As mentioned, data sharing to GISAID^41^ has been beneficial to the wider scientific community and in particular, public health. Therefore, to support users who were already sharing data to GISAID to also share publicly with the INSDC, the GISAID spreadsheet converter was developed and released in November 2021. Note, in addition to this tool, the ENA enables users to reference their corresponding GISAID submission to a sample submission at the ENA, via inclusion of a user-defined field of ‘GISAID Accession ID’.

Utilising this tool, any spreadsheet that was used to submit sequence metadata to GISAID can be converted into the corresponding sample XML which would feed into the ENA’s submission process. This is achieved by mapping metadata fields between the two repositories, reducing burden on the submitter. The tool can be accessed on GitHub^42^.

###### ENA Bulk Webin-CLI Tool

Webin-CLI, a Java based submission tool developed and maintained by the ENA, enables users to pre-validate their submission to identify any potential errors prior to submission. This tool often requires users to develop scripts to suit their submission, along with other preparatory steps for the submission process. This creates a bottleneck when submitting in bulk. The ENA Bulk Webin-CLI tool^43^, fully released in September 2021, is a wrapper tool on top of Webin-CLI which helps to simplify the submission process for users. This tool requires users to prepare a spreadsheet of data to submit, which is provided to it in order to coordinate the submission on behalf of the submitter. This includes automated generation of manifest files, along with automated submission. Prior to usage, users are advised to familiarise themselves with the ENA submissions documentation. Although developed to support COVID-19 submitters and utilised in the backend for the SARS-CoV-2 drag and drop uploader tool, this tool was made available for all ENA submitters for any of the data types that can be submitted through Webin-CLI.

###### ENA Analysis Submitter

As part of the data hubs system, raw read datasets were analysed, however could not be archived at the same time. This contributed to a lag in the number of analysed datasets, compared to datasets being shared. To help reduce this backlog, automation of this step and running this step on-the-fly, was required. The ENA Analysis Submitter^44^ was developed and launched in March 2022. Although required as part of a wider system (in the data hubs), it was developed with external users in mind. Therefore, the tool takes several parameters appropriate for the user’s submission and generates corresponding XMLs for submission via Webin REST API. The tool also uploads data files and carries out the submission process on behalf of the user. As mentioned, it is currently utilised within the ENA Pathogen Analysis System (PAS), described below, to automatically archive resulting analyses back to the data hubs.

##### Data Ownership and Attribution

When sharing data publicly, referencing ownership and provenance was a key concern to collaborators. Despite ownership already being stated within data and metadata records, we felt that more could be done to support researchers in open data sharing. To ease concerns and reduce the barrier to submissions, EMBL-EBI collaborates with ORCiD^45^ to enable researchers/curators to add previously deposited datasets to their ORCiD profile, obtaining credit attribution for data records to which they have contributed. This tool allows you to retrospectively claim datasets to your ORCiD profile^46^.

In addition to this, users can contact the helpdesk, where EMBL-EBI can issue Digital Object Identifiers (DOIs), traditionally used for publications^47^, for their datasets. This is enabled via the BioStudies repository at EMBL-EBI^48^, where a user can register a BioStudy, link their project registered at the ENA, a publication and/or dataset held in another repository to this BioStudy, and then obtain a DOI on the BioStudy.

Claiming datasets through ORCiDs and DOI issuing are new ways for people to obtain credit for sharing data to an open repository and is anticipated to encourage open data sharing in the future. At the time of writing, the number of datasets shared at EMBL-EBI repositories that have been claimed through the ORCiD data claiming system is 2,783.

##### Data Mobilisation

Deposition of the initial Wuhan strain within the International Nucleotide Sequence Database Collaboration (INSDC) - MN908947^49^, collected in December 2019, occurred in early January 2020. Subsequently, the health authorities of China and beyond, demanded further data sharing through GISAID, which then resulted in rapid sharing of SARS-CoV-2 genomes. However, data shared is not visible to all, and therefore not entirely open. To address this, an initial step in facilitating the public sharing of open data was the establishment of a specific support team for submitting centres to share COVID-19 data to the COVID-19 Data Portal. This support was largely provided in the form of detailed documentation, tracking of issues that data submitters faced, meetings with submitting institutes, groups and consortia.

Data sharing within the platform is represented by exponential-looking curves, shown in Figure 3. Here there was a large increase in data sharing for both genomes and raw reads around the Summer months of 2021, which continued into 2022, reflecting the Delta and Omicron variant waves^50^. By the end of Summer 2022, the data portal had seen >6.3million genomes and >5.5million raw reads shared, and since then the rate of data sharing has slowed down. At the time of writing, there have been >6.6million genomes and >6.2million raw reads shared. Tools developed have supported data sharing, however it is not clear if they were all significantly used from Figure 3 alone, there were likely other reasons for the growth in data sharing, such as increased genomic sequencing in certain time periods for surveillance. Observed data sharing aligned relatively closely to sequencing efforts within countries sharing data. The median time between sample collection and dataset sharing was 24 days (Fig.4).

**Figure 3).**
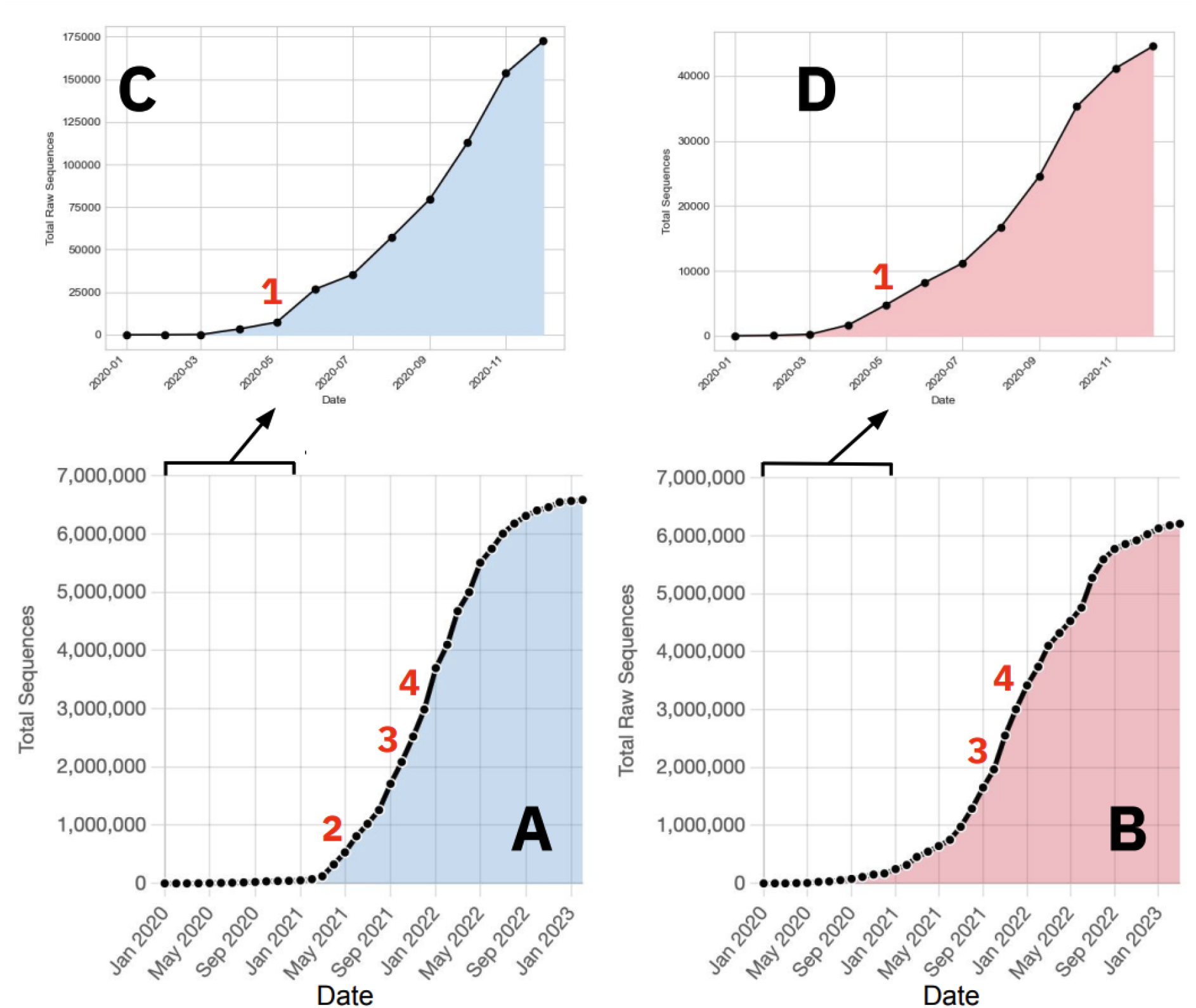
Data mobilisation for (A) nucleotide sequences (genomes) and (B) raw reads from January 2020 up until February 2023. A zoom-in for 2020 has been shown for genome (C) and raw read (D) data mobilisation. Graphs have been annotated with release dates for submission tools. 1 - Drag and Drop Uploader; 2 - Webin SARS-CoV-2 Genome API; 3 - ENA Bulk Webin-CLI tool; 4 - GISAID Spreadsheet Converter tool.

**Figure 4).**
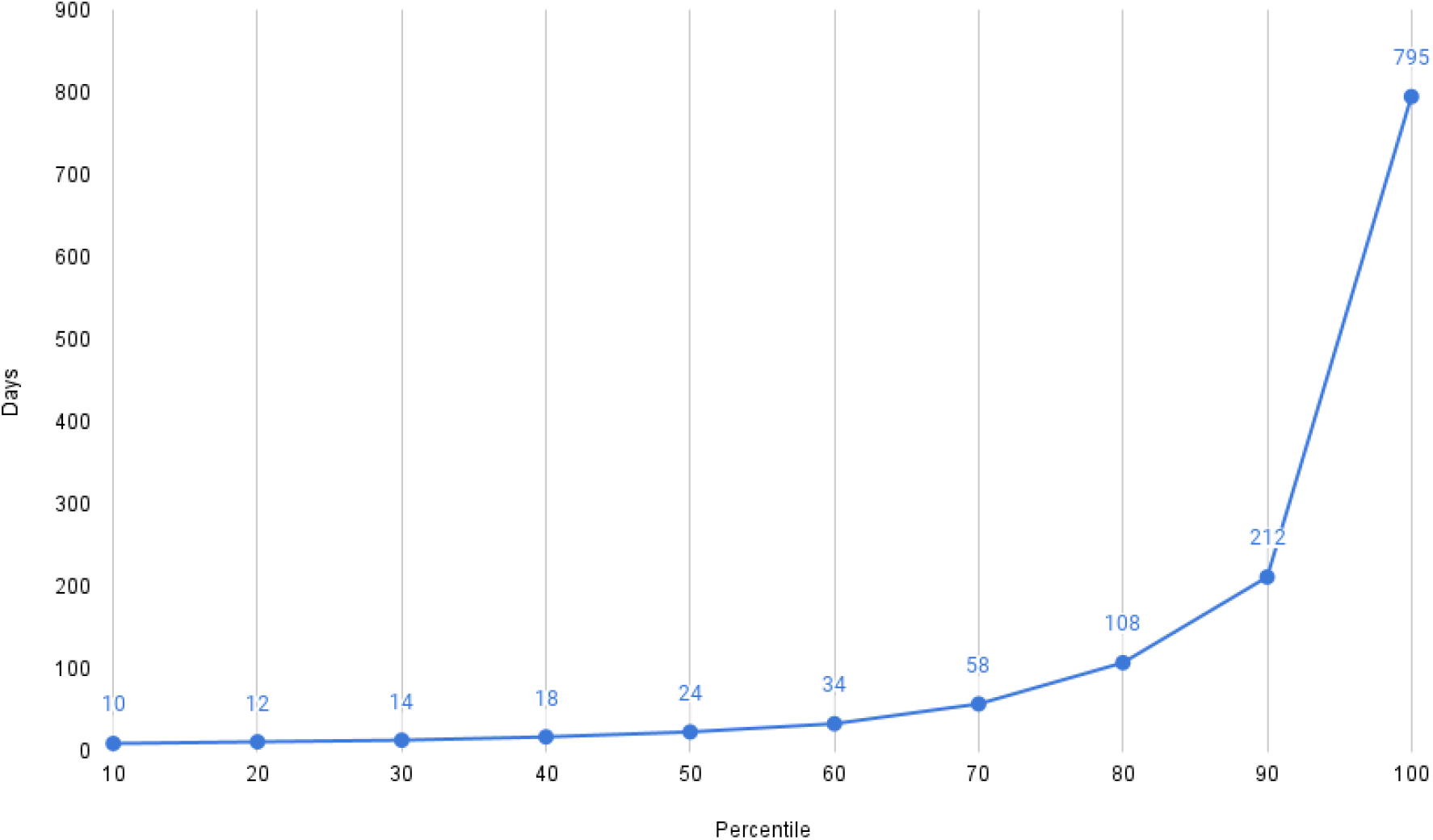
Days from collection to first public - A plot showing the percentile distribution of the length of time from collection date of a sample as reported by the user, to the read data going public in ENA. A median time from collection to public of 24 days is observed.

Geographically, 120 and 103 countries have shared genomes and raw reads, respectively (Fig.5). By 2020, genomes from 75 countries and raw reads from 54 countries had been shared, and by 2021, data had been shared by the majority of countries - 108 for genomes and 96 for raw reads. A summary of the number of datasets shared can be found on the COVID-19 Data Portal, under Data Statistics^51^.

**Figure 5).**
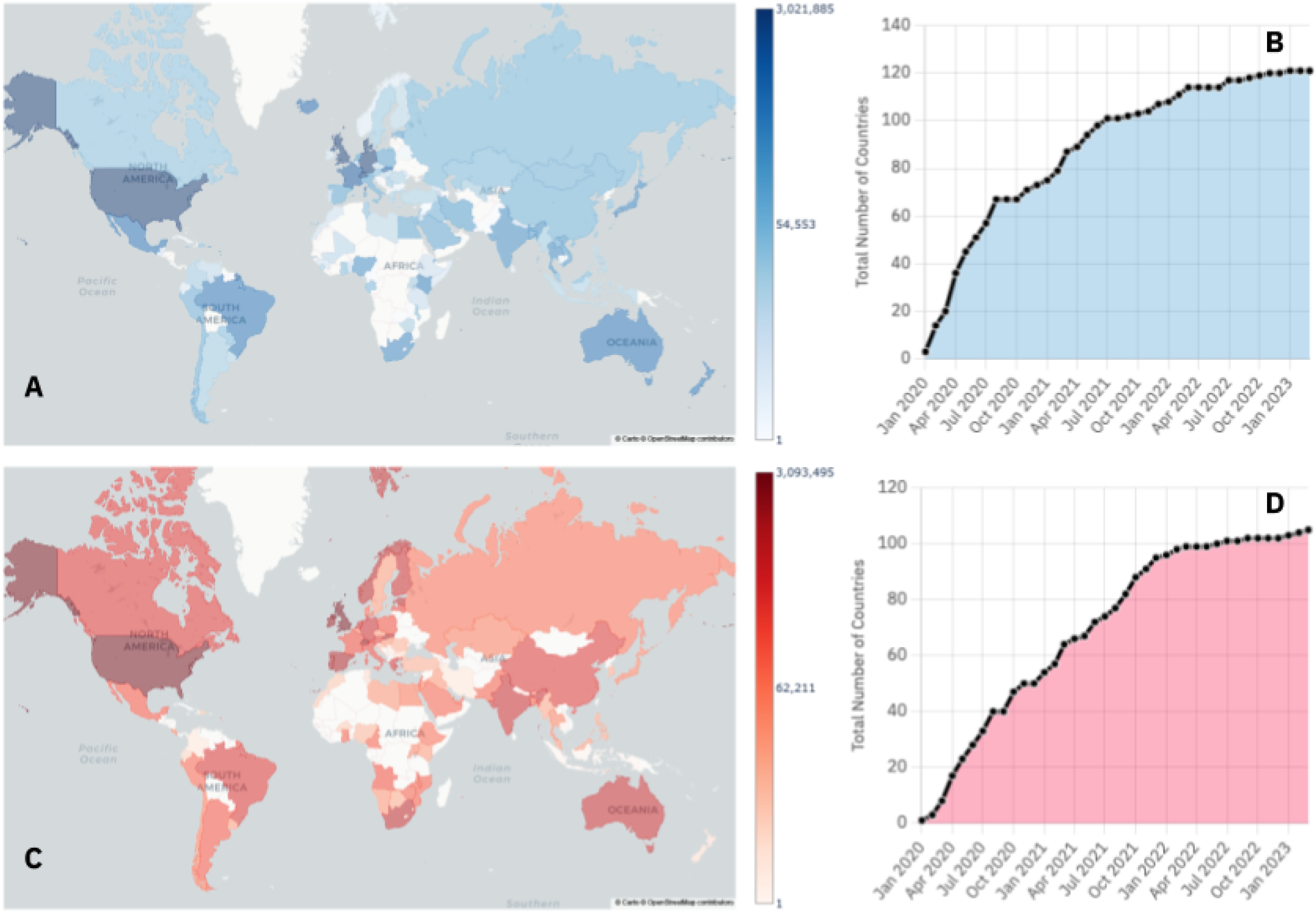
Distributions of data sharing for SARS-CoV-2 genomes (A and B) and raw sequencing data (C and D) from January 2020, until February 2023. The maps present the geographic spread of data sharing, the deeper and darker the colour, the higher the amount of genomes (A) or raw reads (C) shared. The plots represent the number of countries worldwide that have shared genomes (B) and raw reads (D).

**Table 2).** Number of SARS-CoV-2 raw read and genome submisions from January 2020 until March 2023, using the various submission tools. N/A reflects the fact that certain types of submissions cannot be carried out by specific tools. * Represents existing submission routes prior to COVID-19 pandemic.

| Submission Tool | Raw Read Submissions | Genome Submissions |
| --- | --- | --- |
| Drag and Drop Uploader | 5,264 | 1,658 |
| Webin Submissions Portal (Interactive)* | 110,198 | N/A |
| Webin-CLI* | 182,393 | 636,850 |
| Webin SARS-CoV-2 Genome Submission API | N/A | 2,858,579 |
| Webin-REST (Programmatic)* | 2,823,226 | N/A |

### Systematic Analysis

Broadly, the flow depicted in Figure 6 was achieved, where public read data was processed via a reference-based mapping workflow. The pipelines call variants, generating an unfiltered and filtered VCF, and a fasta consensus sequence, per run processed. These analysis products are archived in the ENA and ingested by services and tools, including presentation services such as the ENA browser, Pathogens Portal^52^ and COVID-19 Data Portal. Additionally, data is ingested by visualisation tools, such as the COVID-19 Phylogeny and CoVEO variant explorer. To ensure appropriate tracking and staging of data processing and automated analysis, the ENA Pathogen Analysis System (PAS) ensures that pipelines are processing and archiving data and maintains a provenance trail from input to output, including technologies that utilise cloud compute.

**Figure 6).**
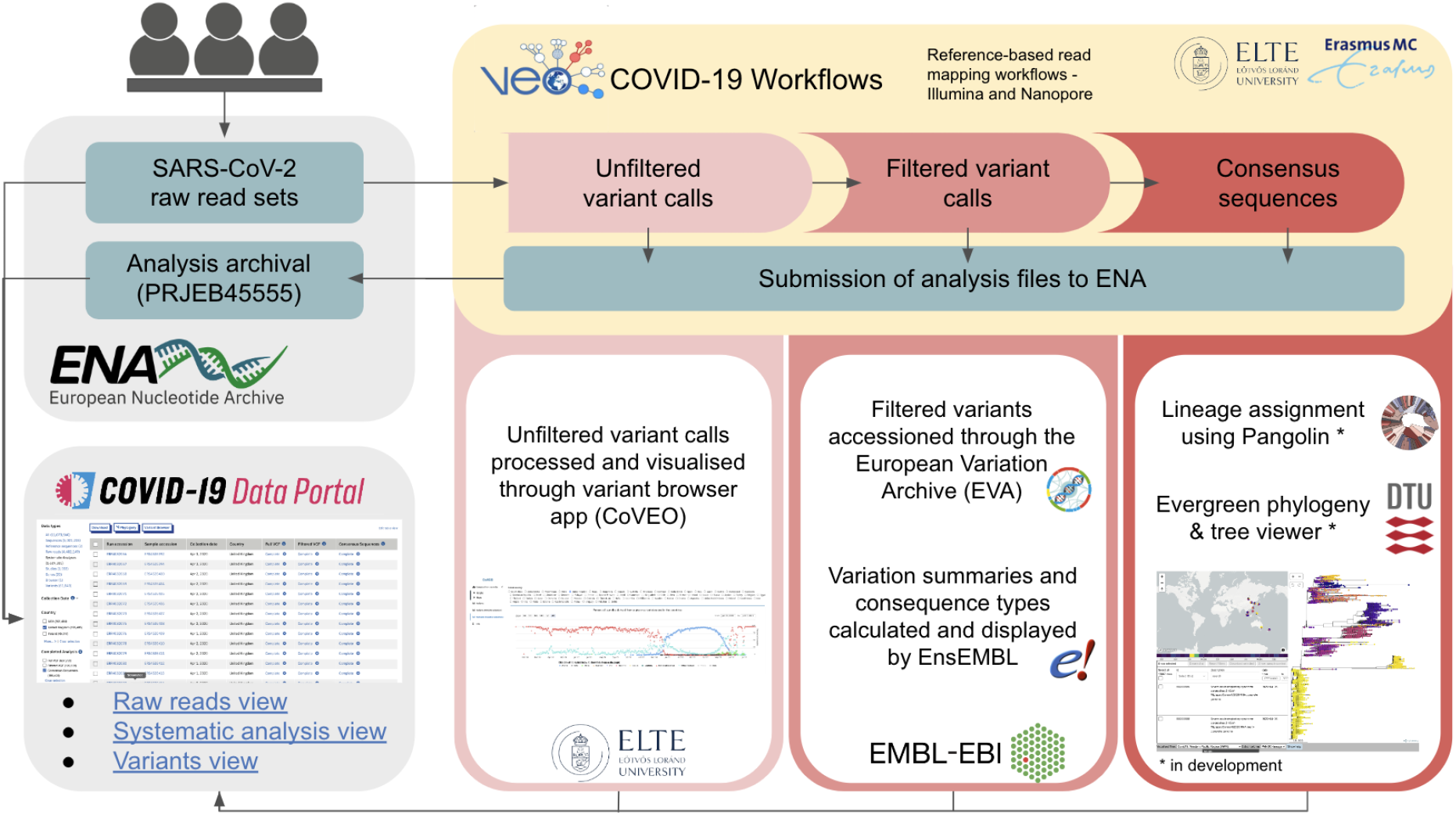
Schematic representing an overview of the systematic analysis of public raw read data shared with INSDC and the COVID-19 Data Portal. Analysis outputs are ingested by various tools and services. Unfiltered variant calls, ingested directly by CoVEO, filtered variant calls by EVA and Ensembl and consensus sequences ingested by Pangolin.

As of writing this paper, 6,239,878 raw read datasets (sequenced using Illumina and Oxford Nanopore technologies) have been shared with the COVID-19 Data Portal, 3,639,538 raw read datasets (Fig.7) have been successfully processed, equating to 58% of the entire raw dataset. The ENA Analysis Submitter tool supported the increase in processing after being incorporated into the ENA PAS to automatically submit data following analysis (Fig.7).

**Figure 7).**
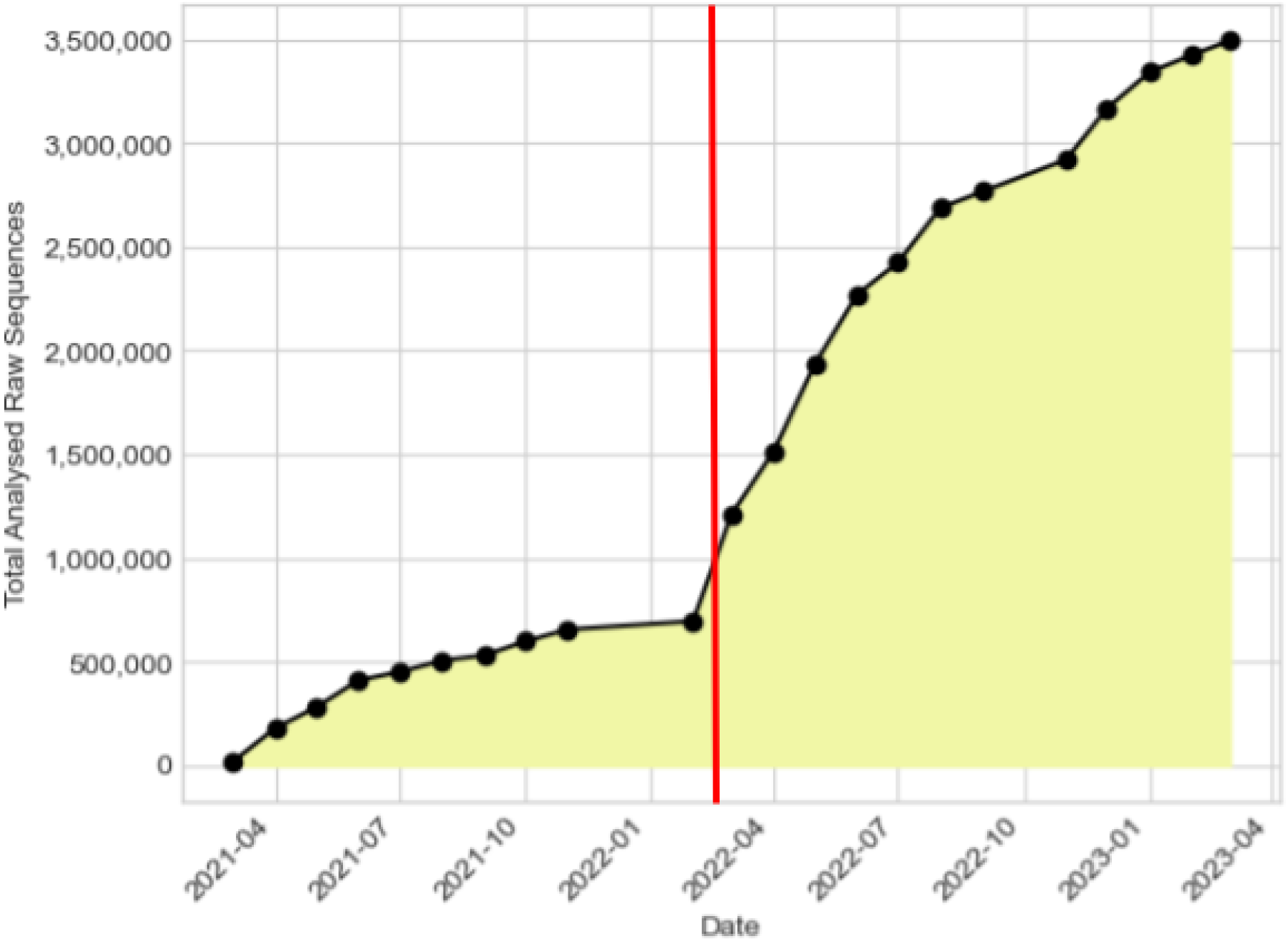
Number of analyses archived following systematic analysis, since March 2021 to March 2023. Generated by tracking the contents of analysis projects detailed in Data Availability. The red vertical line represents the full release of the ENA Analysis Submitter and integration into the ENA PAS.

### Lineages

The COVID-19 Data Portal presents Pango lineage and (where necessary) WHO variant annotations for user-submitted^53^ genomes and for systematically analysed (generated)^54^ genomes. The integrated Pangolin workflow runs three times per week on EMBL-EBI’s high performance computing (HPC) cluster infrastructure. This running frequency was required due to the sheer volume of data submissions on a weekly basis. The resulting lineage assignments are fed back and presented within the COVID-19 Data Portal, which enables for querying and searchability. The workflow currently runs Pangolin for all sequences each time, but a move towards a more incremental approach is likely, only calling lineages for new sequences.

Figures 8 and 9 show that on the whole, submitted sequence data correlated to the raw data being systematically analysed, depicted by similar percentages of WHO and Pango lineage annotations. Along with lineage annotations, the portal also presents a table of submitted genomes that are representative of individual Pango lineages, the ‘representative sequences’^27^.

**Figure 8).**
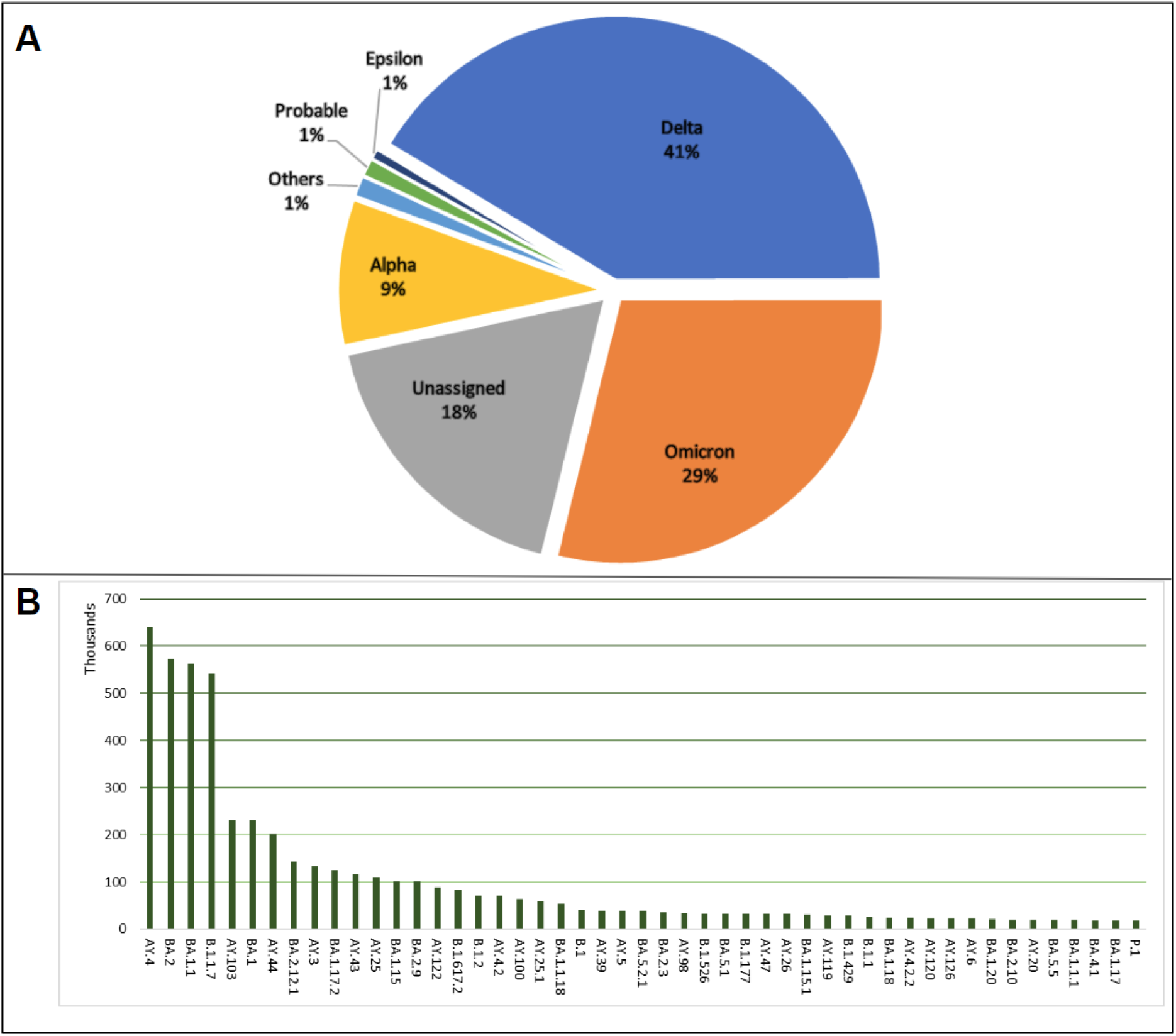
Lineage distributions for SARS-CoV-2 submitted sequences, as of March 2023, A) WHO lineages, Others: consist of a collection of lineages with distribution under 1% which include A.23.1-like, A.23.1-like+E484K, AV.1-like, B.1.1.318-like, B.1.1.7-like+E484K, B.1.617.1-like, B.1.617.3-like, Beta, Eta, Gamma, Lambda, Mu, Theta, Zeta, Iota. B) Top 50 Pangolin lineages observed amongst the dataset.

**Figure 9).**
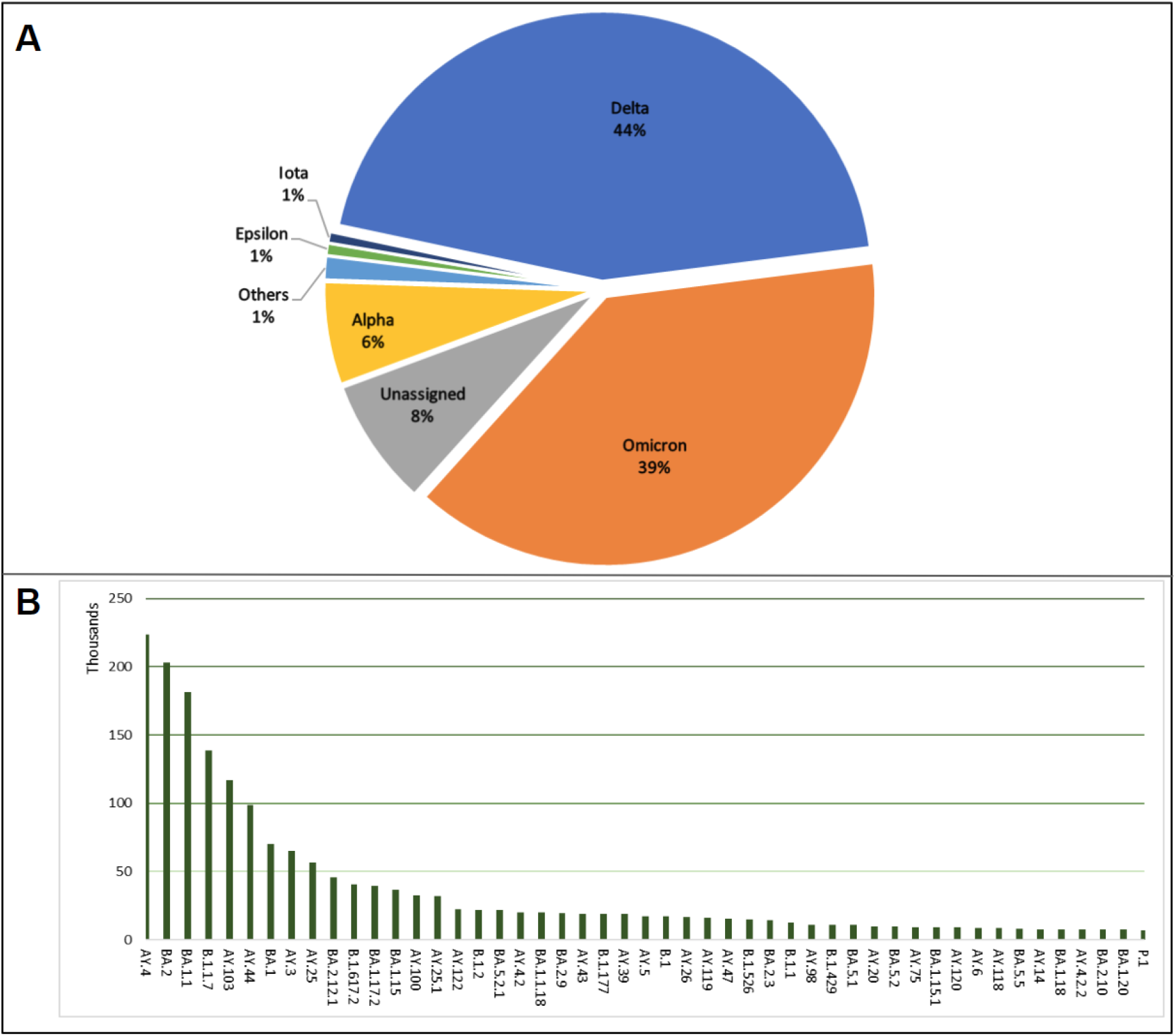
Lineage distributions for consensus genomes generated from SARS-CoV-2 systematic analysis, as of March 2023, A) WHO lineages, Others: consist of a collection of lineages with distribution under 1% which include A.23.1-like, A.23.1-like+E484K, AV.1-like, B.1.1.318-like, B.1.1.7-like+E484K, B.1.617.1-like, B.1.617.3-like, Beta, Eta, Gamma, Lambda, Mu, Theta, Zeta and Probable Omicron. B) Top 50 Pangolin lineages observed amongst the dataset.

### ENA Pathogen Analysis System

The scale of the data to be analysed required a processing management system, to ensure data was not analysed more than once, whilst maintaining the integration of workflows. Therefore, the ENA PAS was developed to support management and scheduling of data processing (Fig.10). It includes several tools and technologies, described below, that have been combined to retrieve information on datasets to analyse, provide infrastructure for analysis through integrated workflows (compute and storage), schedule dataset processing, run analysis, automatically submit resulting analyses, log processing and track overall processing.

**Figure 10).**
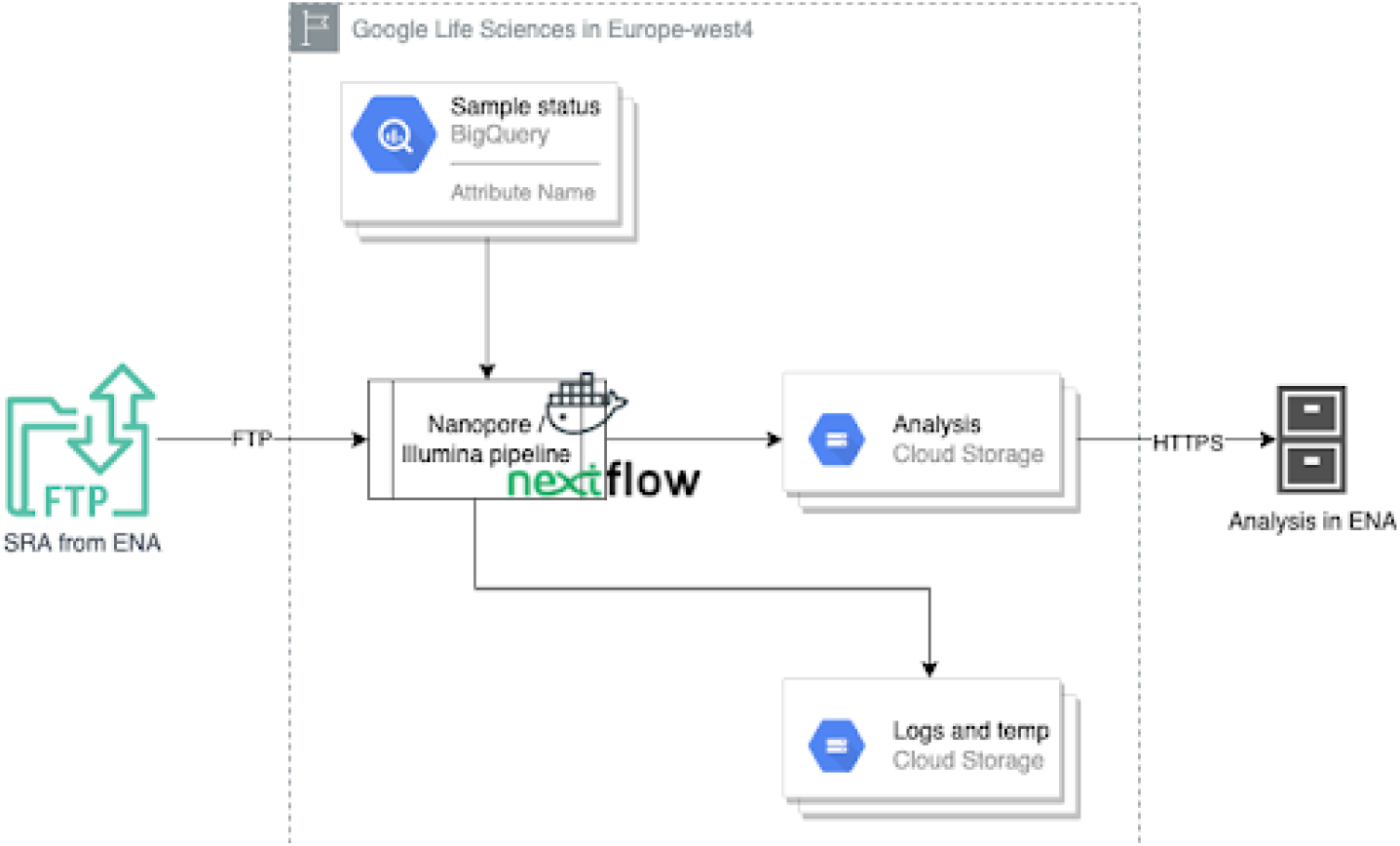
Diagram presenting the flow of data within the ENA Pathogen Analysis System, including Google Cloud services - BigQuery and Cloud Storage. Datasets are pulled using File Transfer Protocol (FTP) and processed using Nextflow pipelines and Docker images, with resulting analyses submitted back to the ENA and the Data Hub.

**Figure 11).**
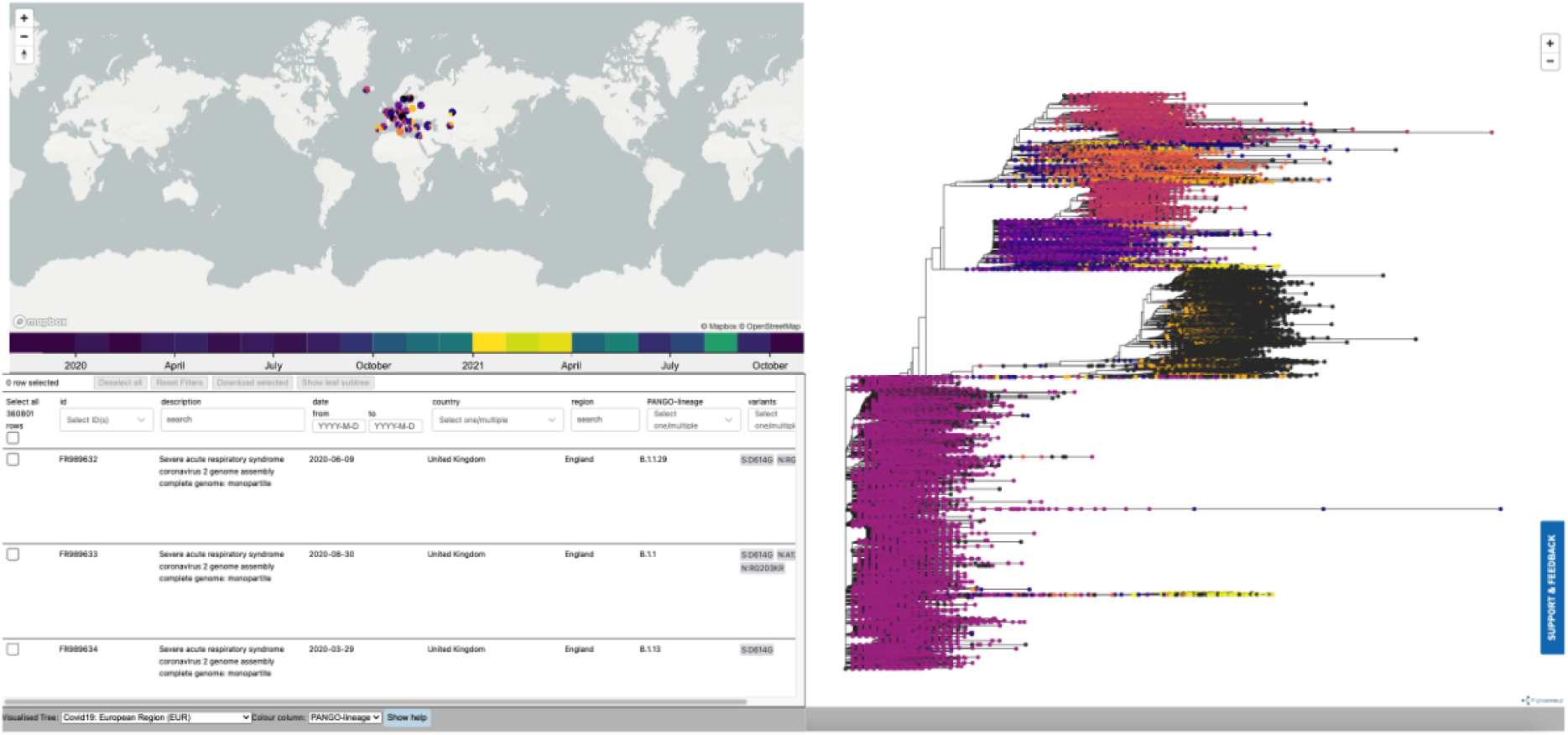
COVID-19 Phylogeny, as of February 2023, presenting in the COVID-19 Data Portal. This includes three main panels - a map on the top left, metadata table bottom left, and phylogenetic tree on the right.

**Figure 12).**
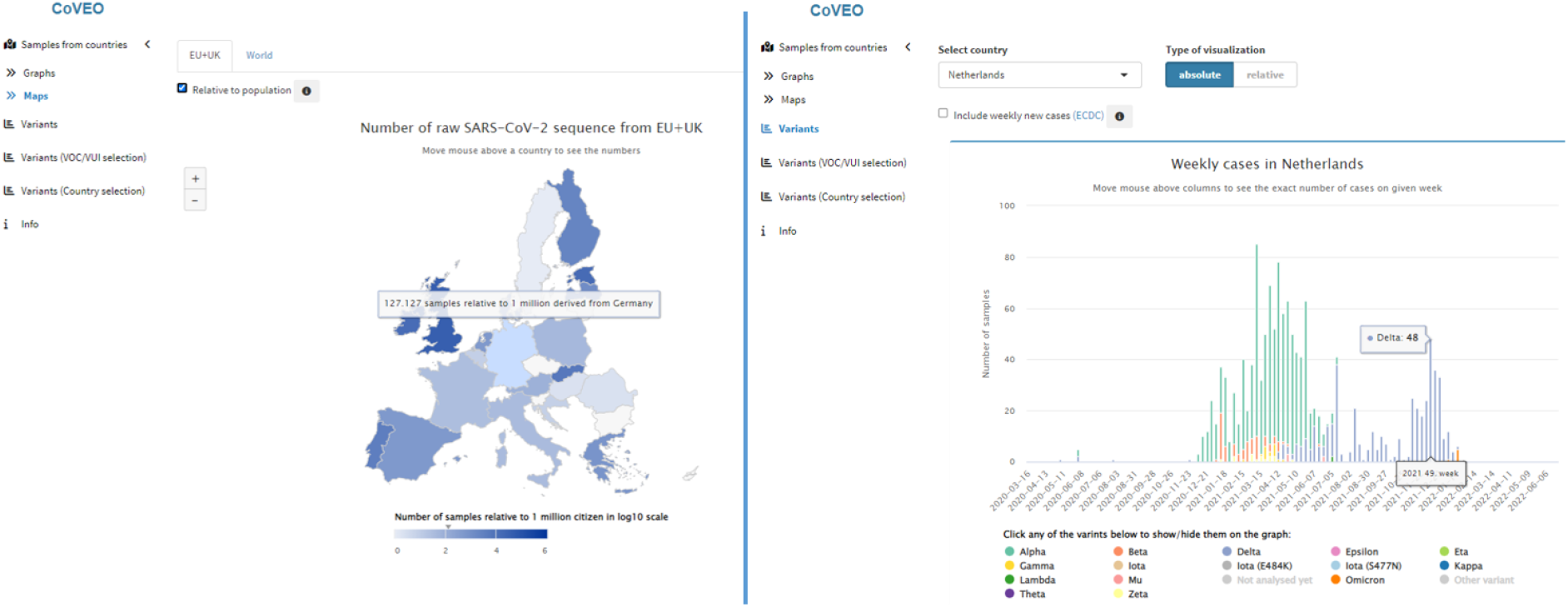
CoVEO variant browser - presenting visualisations, as of February 2023, around the genomic data that is systematically analysed, in the context of VOCs and VOIs.

Nextflow is a workflow management tool used here for scheduling and processing datasets in a parallel and compartmentalised manner^55^. It enables fine-grained control of compute requirements and provides logging capabilities. The workflows are containerised using Docker^56^ for Google Cloud Compute and Singularity^57^ for High-Performance Clusters. Input datasets are streamed just-in-time for processing and results are submitted on-the-fly using the ENA Analysis Submitter tool, described above. A Google BigQuery^58^ database is used to track the number of analysed datasets and submitted results.

### Archival and Data Availability

#### Analysis Archival

The COVID-19 Sequence Analysis Workflow for Illumina and Nanopore data generates three products for each run processed: (1) an archive containing an unfiltered VCF file, a bam file, and a coverage file; (2) a filtered VCF file; and (3) a consensus sequence. These outputs are stored in individual projects under the umbrella project PRJEB45555 to make it easy for users and downstream services to find and retrieve them, while also preserving metadata and raw data links through sample and read references.

To facilitate this archiving, two new analysis types were created at the ENA: COVID19_CONSENSUS, which is used for the computed consensus sequences (output 3); COVID19_FILTERED_VCF, which corresponds to the computed filtered VCFs (output 2); and re-use of PATHOGEN_ANALYSIS, which is used for the archive containing the unfiltered VCF file, bam file, and coverage file (output 1).

For additional discoverability, a view of these analysis products can be found on the COVID-19 Data Portal. This provides the user with more advanced search and download capabilities.

#### Variant Archival and Availability

The systematically generated filtered VCFs feed into a processing flow developed by the European Variation Archive (EVA)^59^, which ingests and accessions the individual variants providing a globally unique identifier (rsIDs) for each one. These accessioned variants have been indexed and displayed on the COVID-19 Data Portal, providing a concise view of all variants, with advanced filtering options. Users can filter on variant types, or use the genome browser to filter on a specific gene or region of interest. The variants are also available to retrieve from the EVA portal via studies PRJEB45554, PRJEB55355 and PRJEB57993. Further downstream, variant calls are fed to Ensembl^60^ and its Variant Effect Predictor (VEP)^61^, to annotate aspects such as consequence types, presenting within the Ensembl COVID-19 browser^62^.

### Visualisation

#### COVID-19 Phylogeny

Using Evergreen, a phylogenetic analysis workflow30 was developed by the VEO consortium and integrated within the data hubs system at EMBL-EBI. PhyloDash enables users to view and search all fields of the metadata table, with the sample metadata optionally supplemented with genotyping data or other phenotypic information, enabling quick selection of samples of interest (Fig.11). Immediate phylogenetic context of a sample can also be viewed by clicking the “Show sample context” button, which provides a subtree containing the given sample.

### CoVEO

CoVEO^36^ was developed by the VEO consortium and integrated within the data hubs system at EMBL-EBI. This includes an underlying PostgreSQL database which was updated with an unfiltered VCF and coverage file per sample processed in the systematically analysed dataset, aligning with the regular monthly VEO bulletin reports.

The CoVEO app presents an interactive visualisation on graphs and maps about the number of raw SARS-CoV-2 sequences submitted by countries across the world and the distribution of variants across time (Fig.12). Users can even input an inclusion/exclusion list of mutations of the S protein, with the app showing the distribution of samples containing the custom-selected mutations across time grouped by country. By enabling focus on samples with sufficient sequencing depth on predefined positions, a unique approach is provided to users. The underlying codebase has been shared on GitHub^63^.

## Discussion

### Open Data Sharing

#### Scale

The scale of open data sharing over the course of the SARS-CoV-2 pandemic is unprecedented64. To consider the numbers of raw read datasets in terms of records (∼25% of all read datasets in ENA), does not even fully do justice to the scale of archived SARS-CoV–2 sequencing data. The publicly archived raw read data in the ENA for SARS-CoV-2 is 2 PetaBases worth of sequencing information.

Contextually, that is enough sequencing data to sequence over 22,000 human genomes from telomere to telomere at a depth of 30X. That is just the data that has been publicly shared and there are far more consensus sequences available in the biodata sphere, so it stands to reason that there must be even more raw read data not archived publicly. By comparison, the bacterial pathogen with the most raw read data shared to the ENA is Salmonella enterica at 313,388 data sets. Other major viral pathogens have low volumes of data sharing, such as Ebola virus (3,099 read data sets) or Zika virus (2,858 read data sets). In fact, if you were to look for all publicly shared raw read data for the WHO priority pathogens65 not including MERS, SARS or their descendants, you’d find 333,087 read data sets. SARS-CoV-2 has 18 times more public read data sets than all of the other WHO priority pathogens combined. The volume of data being generated and submitted publicly to the ENA presented many challenges to overcome in the development of the Data Hubs system. New challenges brought about new tools for submission, search and retrieval, scalable analysis workflows and visualisation tools.

#### Data Sharing

The unprecedented open data sharing mentioned above, does not include the wealth of data that has been shared to GISAID. Openly shared SARS-CoV-2 genomes number around 6.5 million to the INSDC, whereas GISAID currently contains approximately 15 million SARS-CoV-2 genomes^41^. There is likely overlap between the datasets shared, but the extent of this is not known fully. This also highlights that despite the immense and remarkable efforts undertaken and described here, more can be done to truly support open data sharing. Within INSDC, users can share their data, keeping it private, until public release at the time of publication, which appeals to more research focused sequencing efforts. GISAID offered the benefit of being incredibly rapid from submission to availability of genome sequences. Though as we move beyond a rapid data sharing ecosystem, the need for open sharing of both sequences and raw read data becomes the new priority. Through our GISAID conversion tool, we have made it easier for users who have already shared data through GISAID to submit their data to open repositories like the ENA, thus facilitating more users to bring both their sequences and underlying read data into the public sphere. Furthermore, we have made it possible for users to get credit for the data they share openly through ORCiD data claiming and DOI issuing, helping scientists around the world get recognised for their contribution to open biodata. Each development is a small step forward to removing barriers to open data sharing, both practical and social.

The Data Hubs system has driven the development of a whole new set of submission routes for users looking to share data openly in a facile way, such as the Webin SARS-CoV-2 Genome API or the Drag and Drop uploader. Both of these tools are innovations covering two different demographics of users interested in sharing data. The former being targeted more at high volume, programmatically experienced users and the latter targeted at users with little to no knowledge of ENA submission systems.

Researchers now have a vast sea of data available to them, one that is made navigable by the search and retrieval tools powering the COVID-19 Data Portal and ENA. Thanks to the high quality metadata being collected by the majority of SARS-CoV-2 submitters, it is easier than ever for users to find exactly the right data set required to answer their question, either in the immediate present or long into the future. Users can rapidly filter through data based on numerous metadata fields such as country, collection date or in the case of read data, by sequencing platforms or sequencing approaches e.g. WGS or amplicon.

#### Data Dissemination

Dissemination of variants to EVA and Ensembl provides just a couple of examples of the interconnected web of life science resources at EMBL-EBI to which data from the ENA feeds into. As mentioned above, this provides greater context and annotations to sequencing-related datasets, and highlights the importance of the ENA (and other data repositories at EMBL-EBI) to the life sciences community. However, further than EMBL-EBI, datasets (both raw and systematically analysed) feed into downstream tools and services, including Galaxy workflows^66^, CRG Viral Beacon^67^ and CovSPECTRUM^68^ to name a few. This contributes to a network of publicly accessible tools and visualisations that support the scientific community further in analysing and interpreting datasets.

It is worth highlighting that, as the sequencing data presenting within the COVID-19 Data Portal is sourced from the ENA, SARS-CoV-2 sequencing datasets become part of a larger database, that has existed for more than 40 years, with datasets spanning a plethora of different species and taxa. This provides a solid foundation supporting scientific research around targeted projects and resource development, e.g. European COVID-19 Data Platform.

#### Data Analysis and Processing

The volume of data processed through the analysis workflows is immense, with a growing stream of over 6 million raw read datasets publicly available in the ENA. The data hubs system has brought about two significant developments to facilitate a scalable analysis system. Firstly, a dedicated workflow to handle processing of Illumina and Oxford Nanopore datasets, which aim to provide a uniform approach to analysis, and written in Nextflow for cross compatibility. Having the resultant data products archived publicly in the ENA means that there is a systematically analysed, publicly accessible, set of consensus sequences and variants available to researchers around the globe to learn from the SARS-CoV-2 pandemic long into the future. Secondly, we demonstrated a successful deployment across local high-performance computing and distributed cloud platforms to maximise scalability and flexibility in processing. The utility of writing strong Nextflow based pipelines combined with distributed deployment means an elastic analysis management system has been developed, which, when combined with robust pipelines can be rapidly re-deployed to respond to future pandemic scenarios. Further, the utility would be available for more targeted data hubs, whereby specific groups and targeted efforts make use of tools, such as those described here.

The method described in this paper covers analysis of a large and continuously growing dataset, however as discussed under the limitations below, this may come at a cost to accuracy of analysis. Other groups within the community have systematically analysed raw SARS-CoV-2 reads, including Galaxy Europe. Their approach66 offers an equally valuable dataset, by assessing the incoming dataset, more-tailored analyses can be carried out. This results in fewer overall datasets analysed, but greater accuracy in the analysis. Finding an appropriate balance here is a major challenge, however through open sharing of both types of dataset, they can complement one another, as appropriate. What is common to both however, is they are powered by the open sharing of raw read data, and highlights the value of open FAIR data, as neither objective could be achieved without it.

The lineage workflow mentioned above, currently runs Pangolin for all sequences each time, but a move towards a more incremental approach is likely, only calling lineages for new sequences. However, a caveat of the Pangolin classification system is that it remains relatively dynamic, with new decision trees, lineages and sublineages released regularly. Therefore, when a new version of Pangolin is released, the workflow needs to run on the full sequence set again to ensure the latest decision tree is used. Pangolin itself requires quite involved development and curation, and will eventually lose support, meaning running this indefinitely is not sustainable.

## Limitations

### Big Data Processing

The unprecedented scale of data has posed a challenge when it comes to the amount, and in particular, cost of storage and analysis, for example when using commercial-cloud solutions. Increasingly, cloud providers are venturing into the ‘Genomics’ and ‘Life Sciences’ spaces, which provides an opportunity to better-handle big data^69^, however navigating product costs is key, as these appear for all aspects - analysis itself, data storage and data ingress and egress. Hybrid compute solutions (as described in this paper), coupled with cost monitoring, offer a partial solution to this, especially when considering analysis of big datasets.

However, an alternative approach to alleviate processing issues and delays, may have been to consider working with more targeted sequencing efforts, combined with the usage of data hubs, as opposed to a single large public data hub, as described here.

### Generalising Data

By analysing in a systematic way, there are assumptions made on all of the raw read data ingested, for example when parameters are considered for pipelines. It is possible that in some cases, a generic parameter or threshold filters out major variants or fails to identify specific mutations amongst sequences - these are just a couple of examples. However, when analysing big data, it should also be taken into consideration that it is largely not possible to implement individual parameters and specifications in an efficient manner across a dataset. Furthermore, as raw data is openly available, this invites others in the scientific community to (re-)analyse datasets within the larger set to apply specific parameters or utilise other tools.

### Phylogenetic Trees at Scale

The COVID-19 phylogeny aimed to present all appropriate data, filtering out some data according to criteria mentioned in the Methods. However, this became a challenge, as the number of shared datasets rose beyond hundreds of thousands to millions. Overall, this was due to the challenge in presenting a usable phylogeny report, with such a large number of points. As an example, the European tree (which is the largest regional tree), displays >880,000 samples, which present big data challenges in efficiently processing, but also loading and navigating the tree itself.

Generally, this issue existed amongst the community, as how to best present so many samples on a functional phylogenetic tree report. Looking at other openly available tools such as Nextstrain70, reports are restricted to around 4,000 genomes, with subsampling to appropriately represent the global landscape of shared SARS-CoV-2 genomes, balancing this with a smoother user experience. From this, and reflecting on our experience, another alternative model that our taskforce worked towards, and intend to use for future cases, included enabling the user to input a set of samples on top of a baseline tree of ‘representative sequences’. This provides flexibility and a balance in enabling the user to view samples they are interested in, helping to target specific groups of samples (and therefore likely reduce the sample size), whilst maintaining an overall background view of samples across different lineages.

### Looking Forward

As many components of the data hubs system started to mature, there was an increase in training and outreach carried out. Here, it became evident that hands-on training in using the system is important and of great value to users, data providers and data consumers. Embedding this amongst regionally-relevant highlight examples of the importance of open data sharing for life sciences and in particular, infectious diseases, provides wider context to the (further) development of platforms and components described here. Coupled with the data growth mentioned, this enabled us to focus on the user experience, for example what incentives are available for those who share data openly. Outreach also provides a forum to encourage direct feedback from users, which are taken on board and addressed, overall improving systems and components. We intend to continue training and outreach, advertised via platforms such as the News section of the COVID-19 Data Portal, Twitter, EMBL-EBI training pages, amongst others.

Sustainability is a key aspect in setting up platforms and databases. One of the main take-away points from the efforts undertaken and detailed in this paper, was that the true value of the developments made, are in their re-use. The pandemic has provided an opportunity to utilise sustainable practices in developing tools, components and systems which can be re-used in the future. Since the COVID-19 outbreak, tools and components described in this paper have been repurposed for the Monkeypox (MPox) outbreak in May 2022. This provided a better indication of where efforts would have to be focused in repurposing tools for other outbreak scenarios. For example, spinning up an Outbreaks web page within a portal, providing submission instructions and repurposing of the drag and drop uploader were some aspects that were relatively quick. Other aspects, such as setting up systematic analysis of MPox runs took longer, which was expected, as pipeline testing was required to ensure that output was of sufficient quality, especially in identifying known mutations and variants. Therefore, going forward we are confident that tools and components can be repurposed for future potential outbreaks or generally for pathogen pandemic preparedness. The Pathogens Platform52 provides an interface to facilitate and support these efforts.

### Conclusion

The data hubs system represents a fundamental change in how we approach open biodata in response to pandemics. Not only have more tools been developed to enable users to share data quickly, both privately and publicly, but added value can now be offered in exchange for sharing of data in the form of rapid systematic analysis and visualisations of data. This added value can have a significant impact on data submitters who may not have strong bioinformatic backgrounds and feel unable to do much with their data. They can now access a benefit in kind for their data sharing efforts, which will hopefully encourage more users to participate in the open biodata community. Having an adaptable pandemic response system is essential to our collective pandemic security, and the SARS-CoV-2 Data Hubs offer a strong platform on which to build.

## Acknowledgements

The work described in this paper would not have been possible without the many data providers and data brokers who have shared their data openly. Each data provider has been mentioned within sample and run data records that our analyses link to directly (see Data Availability).

This has been a highly collaborative effort, driven by the dedication of those within the taskforce and the authors of this paper - thank you all.

This work has received funding from the European Union’s Horizon 2020 research and innovation programme under grant agreement No. 874735 (VEO), alongside others which have supported the European COVID-19 Data Platform: grant agreement No. 871075 (ELIXIR-CONVERGE), grant agreement No. 824087 (EOSC-Life) and grant agreement No. 825746 (ReCoDID). Furthermore, this work has also received funding from the European Union Horizon project grant agreement No. 101046203 (BY-COVID). This work has been supported by the Member States of the European Molecular Biology Laboratory (EMBL).

